# MECP2 MBD-ID Module: A Unified DNA/RNA Binding Interface Disrupted in Rett Syndrome

**DOI:** 10.64898/2026.02.15.706016

**Authors:** Josiah A. Peter, Thomas A. Weiser, Robert T. Batey, Lee A. Niswander, Deborah S. Wuttke

## Abstract

Rett syndrome neurodevelopmental disorder is caused by mutations in the gene encoding the epigenetic regulator MECP2. While the MECP2 methyl-CpG binding domain (MBD) is well-characterized, the function of the adjacent intervening domain (ID) remains largely understudied. The ID has been described as a distinct RNA-binding region, yet evidence also suggests RNA competitively displaces MECP2 from DNA. Here, we address these conflicting findings by demonstrating the MBD and ID do not function in isolation but as a synergistic functional unit, establishing a new model for MECP2 function. We show the ID significantly enhances affinity of the MBD for methylated DNA by ∼35-fold. Moreover, together these two subdomains form a high-affinity, promiscuous RNA-binding module, with affinity for structured RNAs increased over 1,000-fold compared to the MBD or ID alone. We find binding to RNA precludes binding to DNA, such that the integrated MBD-ID unit explains the competition phenomenon. Analysis of Rett syndrome-associated ID mutations (R167W, K174Q, and R190H) and a therapeutic MiniGene reveals they do not disrupt methyl-DNA binding but instead selectively weaken RNA and non-methylated DNA binding, thereby disrupting the competitive balance between nucleic acid ligands. Our work establishes the MBD-ID module as MECP2’s central nucleic acid interaction hub, whose disruption provides a potential molecular etiology of Rett syndrome due to mutations in the intervening domain.

**GRAPHICAL ABSTRACT:** 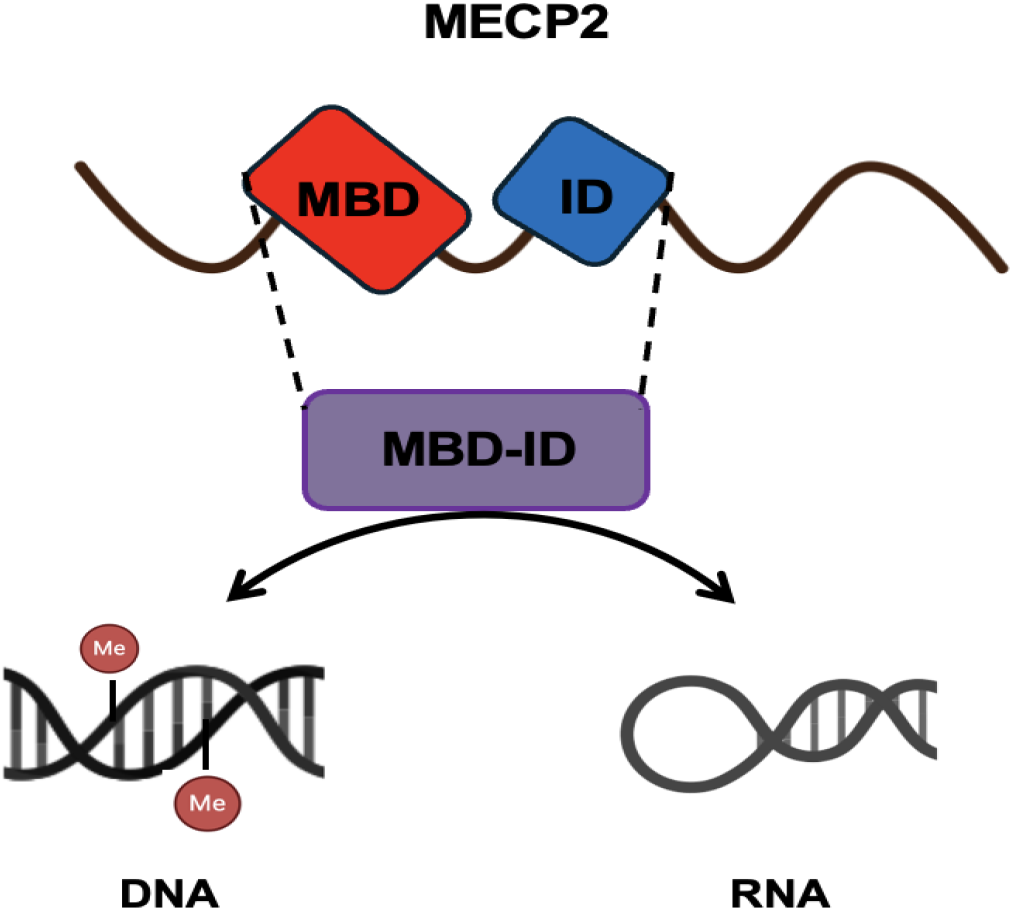

## INTRODUCTION

Methyl-CpG-binding protein 2 (MECP2) is a multifunctional nuclear protein critical for neuronal development and function (1, 2) and mutation of *MECP2* underlies the devastating neurodevelopmental disorder Rett syndrome (1). MECP2 is canonically recognized for binding cytosine methylated DNA and interacting with chromatin modifying proteins (3, 4). As such, MECP2 acts as a master regulator of chromatin architecture (2). MECP2 has broad functional versatility: it influences transcription (5), modulates splicing (6) and miRNA processing (7, 8), and is implicated in the DNA damage response (9), suggesting a multifaceted role in biology that extends beyond simple transcriptional regulation (5). MECP2 functional multiplicity is reflected in its largely disordered domain architecture (10, 11) (Fig. 1A), which includes a partially structured and well-studied methyl-binding domain (MBD), a transcription repression domain (TRD), a NCor/Smrt interacting domain (NID) and an intervening domain (ID) whose specific contributions to the diverse functions of MECP2 remain largely unknown (2). This broad multifunctionality, however, presents a fundamental challenge: how can a single protein achieve its affinity for many different partners, comprised of DNA and RNA and proteins, within the crowded nuclear environment?

**Figure 1:**
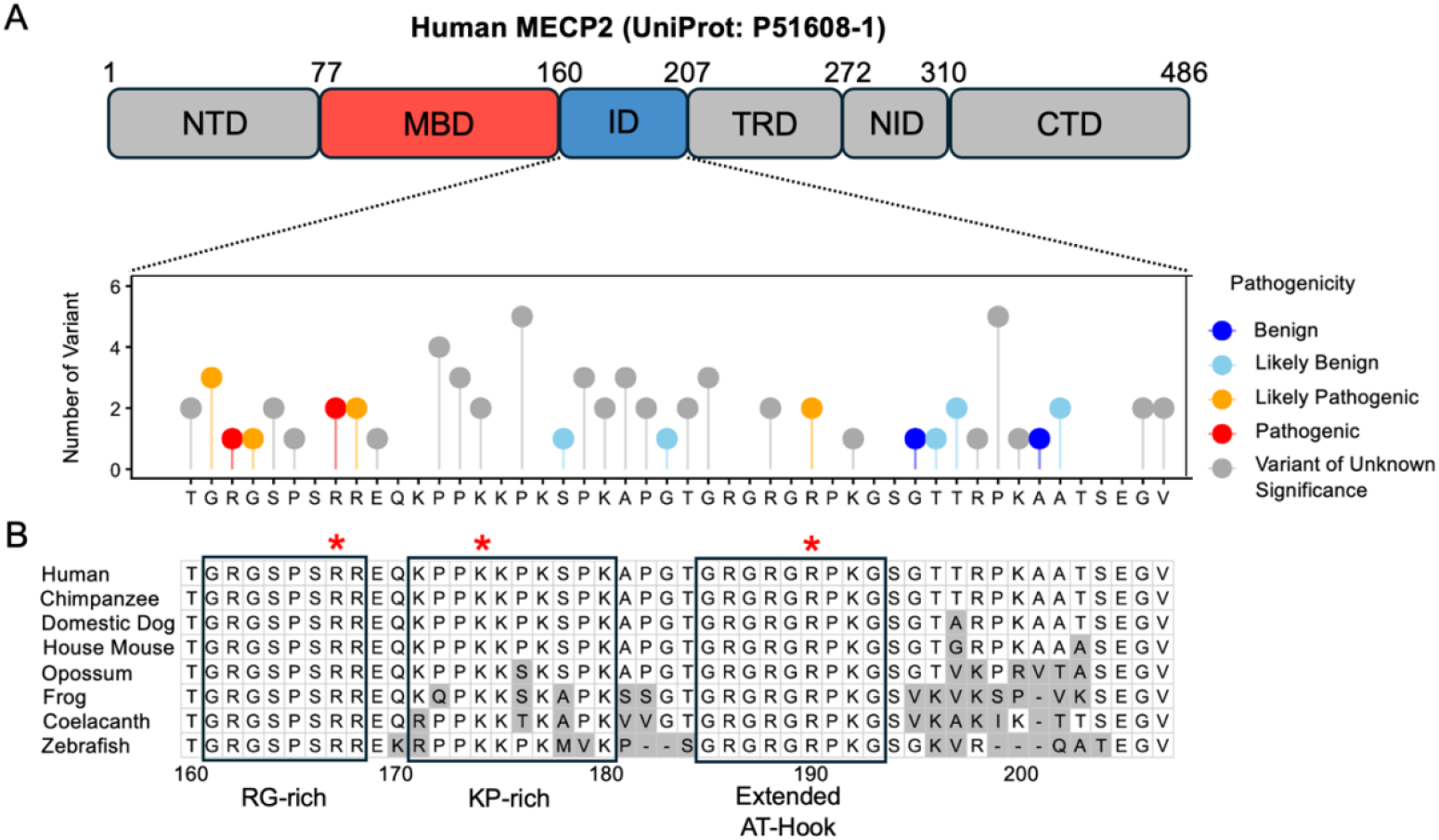
The human MECP2 isoform E2 (UniProt: P51608-1) intervening domain is a conserved region that harbors mutations associated with Rett syndrome and other *MECP2*-related neurodevelopmental disorders. **(A)** Schematic representation of human MECP2 domain architecture. MBD, methyl-CpG binding domain; ID, intervening domain; TRD, transcription repression domain; NID, NCoR interaction domain; NTD and CTD, N- and C-terminal domains, respectively. Below is a lollipop plot of missense variants in the MECP2 ID reported in patients, sourced from ClinVar. The height of each lollipop represents the number of reports for a given residue. Colors denote the consensus pathogenicity classification. **(B)** Multiple sequence alignment of the ID region (human AAs 161-207) across representative species from key vertebrate lineages spanning over 435 million years of evolution. Residues in grey denote evolutionary variation. Disease mutants tested in this study are highlighted with asterisks.

A significant, yet poorly understood, function of MECP2 is its direct interaction with RNA (2, 12). This is underscored by our recent finding that MECP2 directly binds the lncRNA *Rncr3* to control *miR-124a* processing during brain development (13). The current model for the mechanism of RNA binding, based on qualitative assays, proposed a clear segregation of functions: the MBD binds methylated DNA, while the ID is responsible for RNA binding (12). This model, however, conflicts with evidence that RNA can compete with and displace MECP2 from DNA (12). This contradiction implies an uncharacterized functional interplay between the MBD and ID that the segregation model cannot account for. To address this, we sought to determine whether the MBD and ID instead constitute an integrated functional unit governing interaction of MECP2 with both DNA and RNA. Furthermore, we investigated the biophysical consequences of Rett syndrome patient mutations within the ID to assess whether disruption of this potential integrated function could represent a novel disease mechanism.

## MATERIAL AND METHODS

### MECP2 Amino Acid Multiple Sequence Alignment

Multiple sequence alignment was performed using the Clustal Omega algorithm (v.1.2.2) through the Jalview interface (v2.11.5.0) with default parameters, using the full-length sequences from the following UniProt entries: P51608_1 (Homo sapiens); A0A2I3TPC7 (Pan troglodytes); A0A8C0N4P8 (Canis lupus familiaris); Q9Z2D6 (Mus musculus); F6TJ54 (Monodelphis domestica); F6WTL1 (Xenopus tropicalis); H3AT37 (Latimeria chalumnae); A8WIP2 (Danio rerio). The resulting alignment to the ID was visualized using R (v4.4.1) with the ggplot2(v4.0.1) package, with MECP2 missense variants identified and downloaded from the ClinVar database(14) (Human MECP2 protein NM_004992.4; NP_004983 isoform 1, accessed December 2025).

### Cloning, protein expression and purification of MECP2 and MECP2-derived constructs

MBD-ID, MBD, MBD-ID with RTT-Patient ID Mutations (R167W, K174Q, R190H, and K(171-174)A control) and MECP2 MiniGene Human *MECP2* cDNA (Isoform E2, UniProt: P51608_1) and *hΔNIC* (MiniGene) were obtained from AddGene (#48078 and #110191, respectively). The coding sequences for the MiniGene and the subdomains (MBD and MBD-ID) were amplified by PCR and cloned into a pET28a expression vector (EMD Biosciences) using Gibson Assembly (NEB E5510). All constructs feature an N-terminal hexahistidine tag followed by a TEV cleavage site. Rett patient ID mutations (R167W, K174Q, R190H, as well as K(171-174)A control) were introduced into the MBD-ID construct using the NEB site-directed mutagenesis kit (E0552) with oligonucleotides in Table S3. All plasmid constructs were sequence verified and transformed into *Escherichia coli* Rosetta 2 expression cells and plated on LB agar overnight. Protein expression and purification methods were adapted from established protocols (15, 16). For each construct, a single colony was grown in LB media with 50 μg/mL kanamycin overnight at 37 °C. The next day, 1:100 of the overnight culture was inoculated into 1 L 2xYT rich media with 50 μg/mL kanamycin and cultured at 37 °C to an OD_600_ 0.4 - 0.7 and cold shocked on ice for 30 min. Isopropyl β-D-thiogalactopyranoside (IPTG, 0.5 mM final concentration) was added to induce protein expression and cultures were grown at 37 °C for 3 hours while shaking. Cells were harvested by centrifugation at 5000 RCF for 10 min at 4 °C (Avanti JXN-26 floor centrifuge) and pellets stored at −20 °C.

Cell pellets from 1 L culture were thawed on ice and resuspended using a Dounce homogenizer in 40 mL lysis buffer (20 mM Tris pH 7.5, 500 mM NaCl, 10 mM imidazole pH 7.5, 5% glycerol and freshly added one EDTA-free protease inhibitor cocktail tablet (Roche) and 1 mM final concentration of DTT). Cells were lysed on ice using a sonicator (QSonica Q500) (60% amplitude, 3 min total ON-time, pulse: 10 s ON/15 s OFF, three times using ½ inch tip) and the lysate cleared by centrifugation (27000 RCF for 30 min at 4 °C). Cleared lysate was filtered using 0.2 μm syringe filter and loaded into HisTrap column (Cytiva; attached to an AKTA pure FPLC) via a 50 mL superloop. After two washes with 20 mM and 50 mM imidazole in buffer containing 20 mM Tris pH 7.5, 500 mM NaCl, and 5% glycerol, the desired protein was eluted with an imidazole gradient in the same buffer. SDS-PAGE-confirmed fractions were then pooled. The pooled eluate was transferred into a 6-8K MWCO dialysis tubing (Spectra/por – Spectrum Labs) with 10 U/mg of TEV protease (Promega V610) added to remove the hexahistidine tag. Eluate was dialyzed overnight at 4 °C against 1 L of G75 Column Buffer (20 mM Tris pH 7.5, 150 mM NaCl, 5% glycerol, 1 mM DTT). Dialyzed eluate was filtered (0.2 μm filter) and concentrated using 5K and 10K MWCO spin concentrators (Vivaspin Turbo), for MBD and MBD-ID, respectively. Concentrated sample was filtered again before being loaded onto a size-exclusion column (HiLoad 16/600 Superdex 75) and all constructs eluted as a monomer. Pooled fractions containing recombinant protein were assessed for purity using SDS-PAGE, quantified, aliquoted, flash-frozen, and stored at −70°C until use. The MECP2 MiniGene protein is particularly prone to proteolysis; in this case, lysis and wash buffers contained 10 μM leupeptin, 1 μM pepstatin, 0.2 mM phenylmethylsulfonyl fluoride PMSF and 1 μM aprotinin, in addition to EDTA-free protease inhibitor cocktail tablet (Roche). The extinction coefficient used to quantify MBD, MBD-ID, MiniGene, and all MBD-ID-derived constructs is 11460 M^−1^ cm^−1^. A typical yield ranged between 1 – 1.5 mg/L of culture.

#### Full-Length MECP2

The human full-length MECP2 (isoform E2, UniProt: P51608_1) expression vector, containing a TEV-cleavable N-terminal Strep-Tag II, was a gift from Dr. Vignesh Kasinath. The protein expression and purification protocol was adapted from the Kasinath lab with slight modifications. A single transformed colony of *Escherichia coli* Rosetta 2 was grown, IPTG induced, harvested and stored as above. Cell pellets from 1 L cultures were thawed on ice and resuspended using a Dounce homogenizer in 40 mL resuspension buffer (25 mM HEPES pH 8.0, 500 mM NaCl, 10% glycerol, 10 μM leupeptin, 1 μM pepstatin, 0.2 mM PMSF, 1 μM aprotinin, protease inhibitor cocktail tablet (Roche), DNase and salt active nuclease). Cells were lysed and cleared as above. Cleared lysate was filtered using 0.2 μm syringe filter and added into a gravity column, containing a prepared affinity resin (IBA streptactinXT 2-5010-010). The column was sealed and incubated in cold room with gently rocking for 2 hours, followed by 2 washes with 30 mL of low and high salt buffers (0.5 and 1 M NaCl, respectively, in 25 mM HEPES pH 8.0, 10% glycerol, 10 μM leupeptin, 1 μM pepstatin, 0.2 mM PMSF, 1 μM aprotinin, protease inhibitor cocktail tablet (Roche), DNase, and freshly added 1 mM DTT).

After the washes, 15 mL of elution buffer (IBA buffer BXT 2-1042-025) was added into the column. Resin was disturbed and allowed to sit for 10 mins before elute was collected and confirmed by SDS-PAGE. Freshly made ATP and MgCl_2_ (final concentration 10 mM and 20 mM, respectively) was added to the 15 mL elute. The entire solution was transferred into a 6-8K MWCO dialysis tubing (Spectra/por – Spectrum Labs), and dialyzed against 1.5 L dialysis buffer (25 mM HEPES pH 8.0, 300 mM NaCl, 10% glycerol and 1 mM DTT) overnight at 4 °C. Dialyzed solution was filtered and loaded into a heparin column attached to an AKTA Pure FPLC, and eluted with a salt gradient elution buffer (100 mM – 1 M NaCl, in 25 mM HEPES pH 8.0, 10% glycerol and 1 mM DTT) to separate MECP2 from co-purifying contaminants like HSP70. To further polish the protein, the fraction eluted from heparin column was concentrated and loaded onto a size exclusion column (HiLoad 16/600 Superdex 75) and eluted with the sizing buffer (25 mM HEPES pH 8.0, 500 mM NaCl, 10% glycerol and 1 mM DTT), after which high-purity fractions were pooled, quantified, concentrated, aliquoted, and flash-frozen for storage at −70 °C. The extinction coefficient used to quantify the full-length MECP2 is 19940 M^−1^ cm^−1^. A typical yield ranged between 1 – 1.5 mg/L of culture.

#### Maltose Binding Protein (MBP)-tagged ID

As intrinsically disordered domains like the ID often have solubility issues, we cloned the coding region of the ID into a pHMM expression system, a gift from Dr. Robert Batey, that contains a factor Xa cleavable N-terminal MBP, using Gibson Assembly (NEB E5510). The expression plasmid was transformed into *Escherichia coli* Rosetta 2 and a single colony was grown, IPTG induced, harvested and stored as above. Cell pellets from 1 L cultures were thawed on ice and resuspended using a Dounce homogenizer in 40 mL lysis buffer (20 mM Tris pH 7.5, 500 mM NaCl, 10 mM imidazole pH 7.5, 5% glycerol and freshly added one EDTA-free protease inhibitor cocktail tablet (Roche) and 1 mM DTT). Cells were lysed and cleared as above. Cleared lysate was filtered using 0.2 μm syringe filter and loaded into HisTrap column (Cytiva; attached to an AKTA pure FPLC) via a 50 mL superloop. After two washes with 20 mM and 50 mM imidazole in buffer containing 20 mM Tris pH 7.5, 500 mM NaCl, and 5% glycerol, the desired protein was eluted with an imidazole gradient in the same buffer. SDS-PAGE-confirmed fractions were then pooled. The pooled eluate solution was transferred into a 6-8K MWCO dialysis tubing (Spectra/por – Spectrum Labs) and dialyzed overnight at 4 °C against 1 L of G75 Column Buffer (20 mM Tris pH 7.5, 150 mM NaCl, 5% glycerol, 1 mM DTT). Dialyzed eluate was filtered with a 0.2 μm filter and concentrated using 30K MWCO spin concentrators (Vivaspin Turbo). Concentrated sample was filtered again before being loaded onto a size exclusion column (HiLoad 16/600 Superdex 75) and eluted as a monomer. Pooled fractions containing recombinant protein were assessed for purity using SDS-PAGE, quantified, aliquoted, flash-frozen, and stored at −70 °C. To isolate the MBP for testing for nucleic acid interaction (Fig. S3), purified MBP-ID was incubated with factor Xa protease (NEB P801) at the manufacturer recommended ratio of 1:100 (w/w) protease:substrate. The cleavage reaction was then loaded onto an SEC column and purified as described above. The extinction coefficient used to quantify MBP-ID and MBP is 66350 M^−1^ cm^−1^. A typical yield ranged between 3 - 5 mg/L of culture.

### Oligonucleotide preparation

All ligand sequences used for binding experiments are listed in Table S2. DNA oligonucleotides were ordered from Integrated DNA Technologies (IDT) with standard desalting and HPLC purified. For fluorescently labeled ligands used in anisotropy measurements, the sense strand was synthesized with fluorescein (6-FAM) conjugated to the 5’-end. The labeled and unlabeled complementary strands were annealed by slow cooling (heating at 95 °C for 3 minutes, then slow cooled to room temperature for 3 hours) to promote inter-molecular interaction, at 1 μM in annealing buffer (20 mM Tris pH 7.5, 50 mM NaCl).

Fluorescein (6-FAM) labeled RNA for anisotropy measurements were ordered from Horizon Discovery and IDT with standard desalting and HPLC purified. RNA ligands were refolded for binding assays by snap cooling (95 °C for 3 min, ice for >5 min) to promote intramolecular interaction, at a 1 μM concentration in folding buffer (20 mM Tris pH 7.5, 10 mM NaCl). The labeled and unlabeled complimentary strands used to make double stranded RNA were annealed as described above. All oligonucleotides were verified on a polyacrylamide native gel.

### Fluorescent Anisotropy Binding Assay for K_D,app_ Determination

The apparent binding affinities were determined using previously described methods (17–19). The binding buffer (20 mM Tris pH 7.5, 150 mM NaCl, 5% glycerol and 0.025% NP-40) was used to dilute purified proteins and oligos to desired concentration. The full-length MECP2 was stored in 500 mM NaCl and usually first diluted with nuclease free water to bring NaCl concentration to 150 mM prior to serial dilution. The 6-FAM labeled DNA and RNA ligand final binding experiment concentrations were 1 nM and 2 nM, respectively. Purified proteins were serially diluted (1:2 for each dilution) from 10 µM to 2.4 pM in binding buffer. Protein and ligand were mixed in a 20 µL binding reaction in a flat-bottom low-flange 384-well black NBS polystyrene plate (Corning) and allowed to incubate for at least 1 hour at room temperature to reach equilibrium before measurements. Parallel (*I*∥) and perpendicular (*I*⊥) fluorescence intensities were measured with ClarioStar Plus FP plate reader (BMG Labtech), and anisotropy for each protein dilution point was calculated using *Anisotropy* = (*I*∥ + *I*⊥)/(*I*∥ + 2 × *I*⊥). Anisotropy values were plotted as a function of the log of protein concentration. The apparent dissociation constant K_D,app_ was determined by fitting the data to the quadratic binding equation

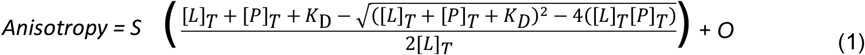

where S is the difference between the maximum and minimum anisotropy values, O is an offset factor equivalent to the minimum anisotropy value, and [P]_T_ and [L]_T_ are known concentrations of MECP2 constructs and DNA/RNA ligands, respectively.

The apparent K_D,app_ of full-length MECP2 vs methylated DNA (mBDNF) was determined by fitting the data to the Hill equation, to determine the K_D_ and Hill coefficient (to access cooperativity). The quadratic and Hill binding equation data fitting were done using an in-house generated python script (https://github.com/nicklammer/AnisotropyBindingFit). Each binding reaction was performed in triplicate with independent dilutions. The statistical significance was determined for differences between average values of K_D,app_ using student’s two-tailed t-test.

Competition experiments were performed under the same fluorescence anisotropy conditions as described above. Here, 1 nM labeled mBDNF promoter was bound at ∼50-70% by the full-length MECP2 (7 nM), MBD-ID (7 nM) and MBD (120 nM) and competed off by titration with unlabeled mBDNF DNA or RNA Rncr3_2. Competitor concentrations were at least 10-fold above and below the K_D,app_. The reaction was equilibrated for at least 2 hours at room temperature. Twelve data points were used per experiment, and experiments were performed in triplicate with independent dilutions. IC_50_ reported as concentration of unlabeled competitor required to reduce the affinity of MECP2 (and sub-domain constructs) to the labeled ligand by 50%. Data were fit using GraphPad Prism10.

### Electrophoretic Mobility Shift Assay (EMSA)

EMSA was performed as previously described(20–22) with modifications. Purified recombinant MBD-ID and MBD were 1:2 serially diluted from 1000-0.5 nM (12 concentration points) in binding buffer (20 mM Tris pH 7.5, 150 mM NaCl, 5% glycerol, 0.025% NP-40). 10 nM of labeled Rncr3_1 RNA was added to the protein (at a 1:1 volume ratio; final volume 20 μL) and allowed to equilibrate for an hour at 4°C. While samples were equilibrating, a 15% acrylamide:bisacrylamide 1X TBE gel was pre-run at 175 V for 15 minutes at 4 °C. The entire binding reaction was then run at 200 V for 30 min at 4 °C. Gels were imaged using Cy2 settings (473 nm excitation) on Typhoon FLA 9500 Imager (GE Healthcare).

## RESULTS

### Molecular Dissection of the MECP2 Intervening Domain (ID)

The MECP2 intervening domain (ID) is an intrinsically disordered (11), 46-amino-acid region (AAs 161-207) that is highly basic and contains motifs associated with nucleic acid binding (20), including arginine-glycine (RG)-rich, lysine-proline (KP)-rich, and extended AT-hook motifs (23). Despite its association with Rett syndrome pathology (14, 20), the molecular function of the ID remains poorly defined. This functional gap is particularly pressing, as the ClinVar (14) database reports numerous missense mutations within this domain associated with *MECP2*-related neurodevelopmental disorders, many of which are classified as variants of unknown significance (14) (Fig. 1A). The strong evolutionary conservation of the ID from humans to zebrafish underscores its critical and non-redundant role (Fig. 1B).

To address this knowledge gap and determine the contribution of the ID to MECP2 function, we adopted a reductionist biochemical approach. We purified recombinant full-length (FL) MECP2 alongside a panel of sub-domain constructs (Fig. 2A) designed to deconvolute the individual and potential integrated functions of the MBD and the ID. These included: 1) the isolated MBD (AAs 78–165), representing the well-studied canonical DNA-binding module(2); 2) the isolated ID (AAs 161–207), implicated in nucleic acid interactions (12, 20); and 3) a construct encompassing both domains (MBD-ID, AAs 78–207). To overcome the solubility challenges of the intrinsically disordered ID, this region was expressed and purified as a fusion with maltose-binding protein (MBP-ID). As a critical control, we confirmed that the MBP tag itself does not interact with nucleic acids (Fig. S3). All proteins were expressed and purified to homogeneity (Fig. 2B).

**Figure 2:**
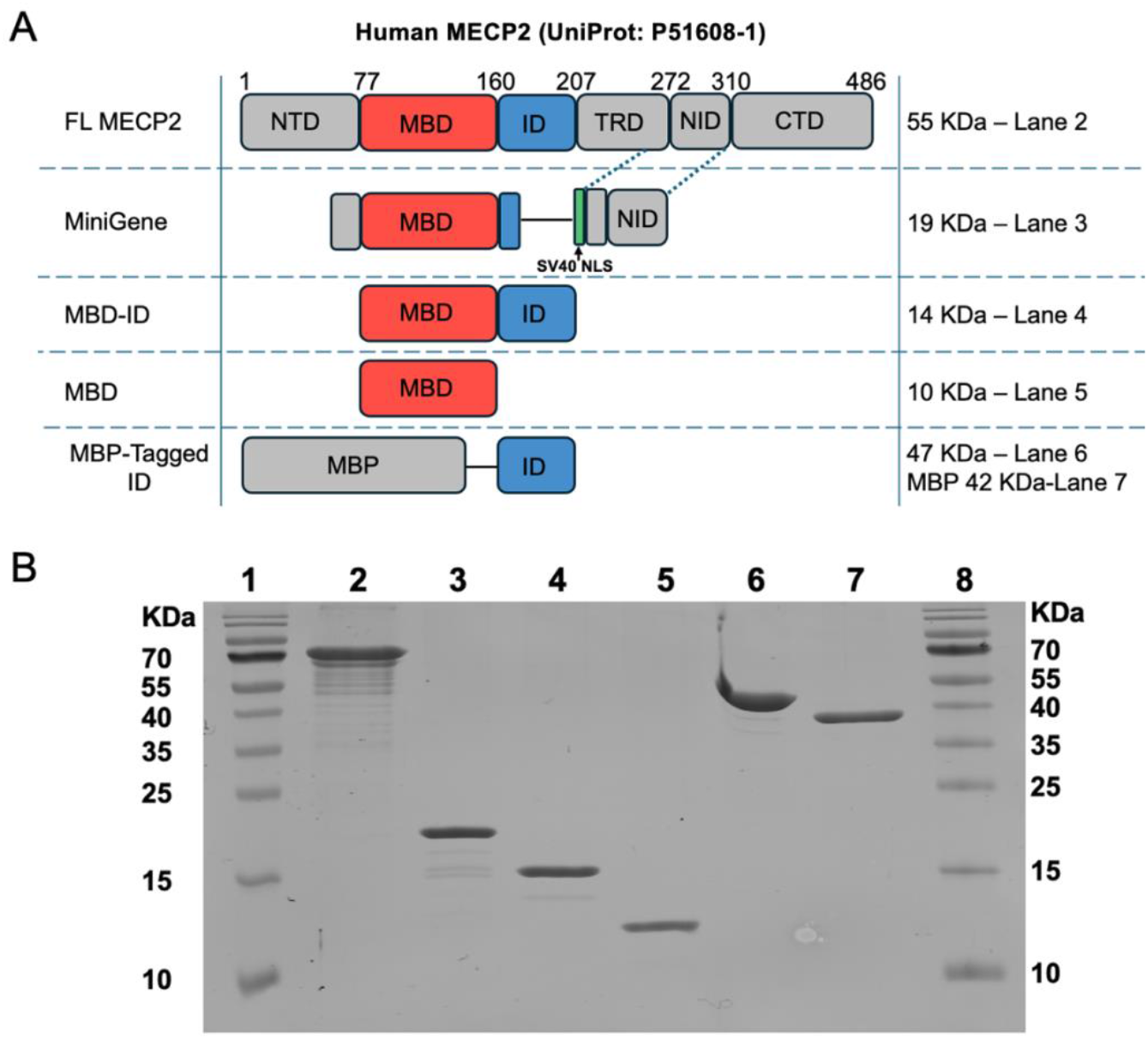
Human MECP2 (UniProt: P51608-1) domain constructs and purified recombinant proteins used. **(A)** Schematic of human MECP2 domain architecture and the derived recombinant constructs used in this study. FL, full-length; MBD, methyl-CpG binding domain; ID, intervening domain; TRD, transcription repression domain; NID, NCoR interaction domain; NLS, nuclear localization signal; MiniGene, therapeutic protein domain structure (hΔNIC). **(B)** AcquaStain-stained SDS-PAGE gel of purified recombinant proteins. MECP2 runs high due to its high proline content.

### Synergy Between the ID and MBD Drives High-Affinity DNA Binding by MECP2

To directly test whether the MBD and ID constitute an integrated functional unit, we first established a quantitative baseline for their individual and combined contributions to DNA binding. We employed an *in vitro* fluorescent anisotropy (FA) assay to determine the affinity and specificity of our purified protein constructs for a standard 15-bp DNA sequence from the BDNF promoter (24), testing both methylated (mBDNF, with a single symmetric CpG methylation) and non-methylated (BDNF) versions. All binding assays were performed in a buffer containing 150 mM NaCl, pH 7.5 at room temperature.

Consistent with prior studies (16, 25–28), FL MECP2 bound mBDNF cooperatively with low nanomolar affinity (K_D,app_ = 2.5 ± 0.1 nM, Hill n = 2.4) and displayed approximately 6-fold tighter binding compared to the non-methylated sequence (Fig. 3A), confirming its canonical function as a methyl-CpG binding protein (3, 4). It is worthy to note that MECP2 has been predicted to harbor a number of molecular recognition features (MORFs) (15) speculated to form secondary and tertiary structure in a methylation density-dependent manner (29), hence the binding cooperativity observed. The FL MECP2 preferential binding for mBDNF over unmethylated BDNF was retained in both the MBD and MBD-ID constructs (Fig. 3B). Strikingly, however, the inclusion of the ID dramatically enhanced the DNA-binding affinity of the MBD by approximately 35-fold (Fig. 3B), consistent with prior observations (15, 30). Moreover, the binding profile of the MBD-ID construct closely recapitulated that of the full-length protein, confirming it is the necessary and sufficient unit for DNA binding by MECP2. In contrast, the ID alone showed weak micromolar affinity to both mBDNF and BDNF with little to no preference for the methylated sequence (Fig. 3B). These data indicate that the ID acts in concert with the MBD to constitute the core DNA-binding module of MECP2. Therefore, the MBD and ID function as an integrated functional unit, with the MBD providing specificity for methylated DNA (24).

**Figure 3:**
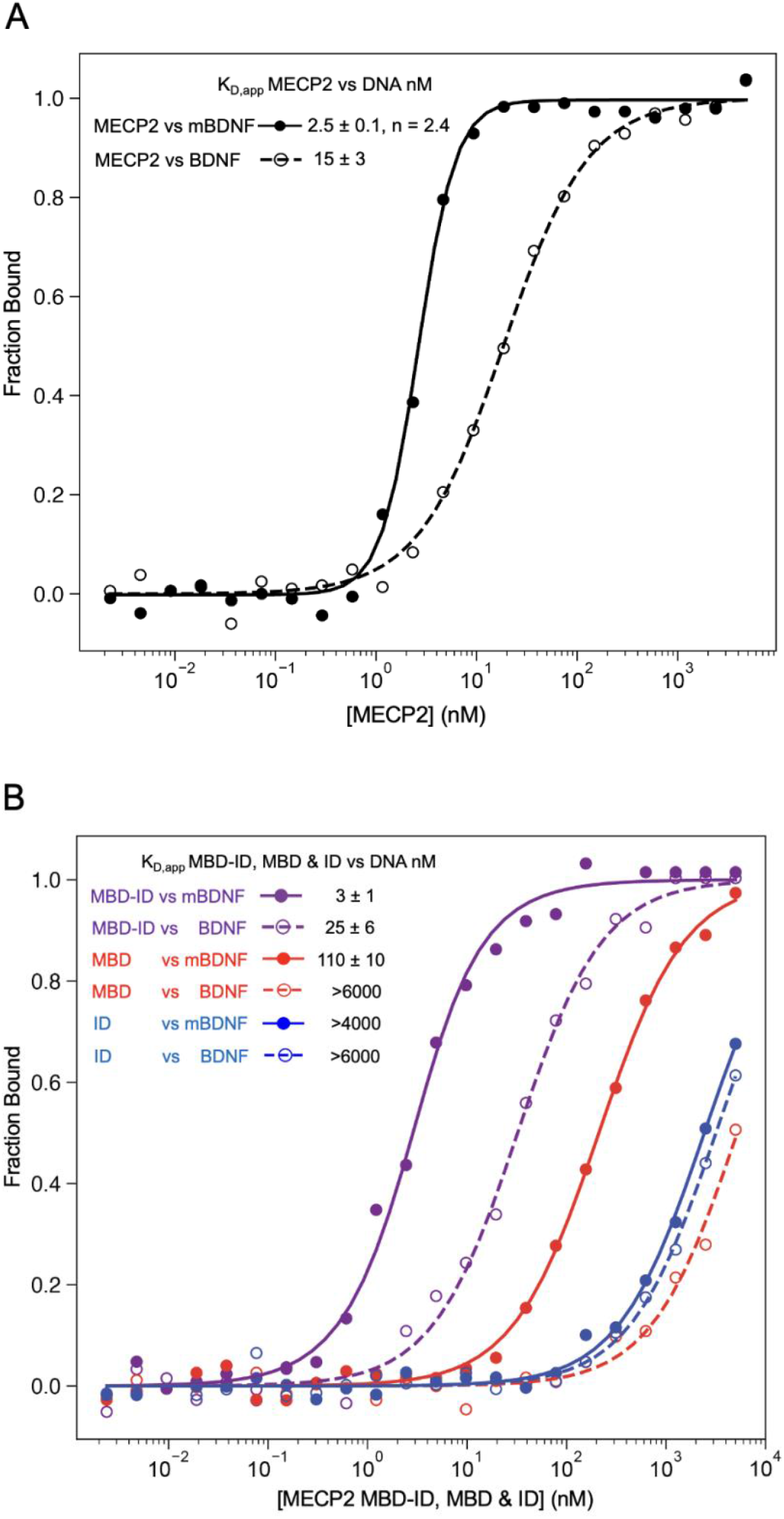
Synergy between the ID and MBD drives high affinity DNA binding by MECP2. **(A)** Fluorescent anisotropy (FA) binding curve for full-length MECP2 bound to methylated and non-methylated BDNF. **(B)** FA binding curve for MBD-ID, MBD and ID bound to methylated and non-methylated BDNF. n>3, standard error of the mean reported.

### The MECP2 MBD-ID is a High-Affinity, Promiscuous RNA-Binding Module

While MECP2 is well known as a methyl-DNA binding protein (2–4, 24, 29), several lines of evidence have suggested that MECP2 can also interact with RNA (2). A 2004 study first proposed that MECP2 possesses distinct DNA- and RNA-binding domains (12), with the ID facilitating RNA interactions potentially due to the presence of basic amino acid patches and an extended AT-hook motif (20, 23) (Fig. 1B). This concept has been reinforced by more recent studies, including RIP-seq experiments that identified a range of MECP2-bound lncRNAs in mouse brain (31) and our recent finding that MECP2 directly interacts with the lncRNA *Rncr3* to control *miR124a* processing during brain development (13). Furthermore, a recent SELEX-based study confirmed that MECP2 can bind a broad spectrum of structured and dsRNAs with high affinity (32), although the role of the ID in this RNA binding was not directly tested.

The quantitative biophysical parameters governing MECP2-RNA interaction and their comparison to MECP2-DNA interactions are not resolved. For example, it is not known if the ID alone is sufficient for high-affinity RNA binding, as initially suggested (12), or if the ID acts in concert with MBD, similar to the results above for DNA interaction. To bridge this gap, we systematically characterized the RNA-binding capacity of the FL MECP2 and subdomain constructs. We found that the FL MECP2 bound a 23-nucleotide hairpin RNA derived from *Rncr3* (Rncr3_1) with high affinity (K_D,app_ = 8 ± 3 nM) (Fig. 4A). Similarly, the MBD-ID construct bound the same RNA ligand with comparable affinity (K_D,app_ = 5 ± 1 nM), as measured by both FA and EMSA (Fig. 4A and B). In stark contrast, the isolated ID or the isolated MBD bound this RNA very weakly (K_D,app_ ∼ 2 µM and >6 µM, respectively; Fig. 4A and B). Hence, the ID alone cannot account for the tight affinity binding to RNA. Instead, the ID in conjunction with the MBD enhances RNA binding by almost three orders of magnitude in comparison to the individual domains, indicating that the MBD-ID is necessary and sufficient to recapitulate the nucleic acid binding behavior of the full-length protein.

**Figure 4:**
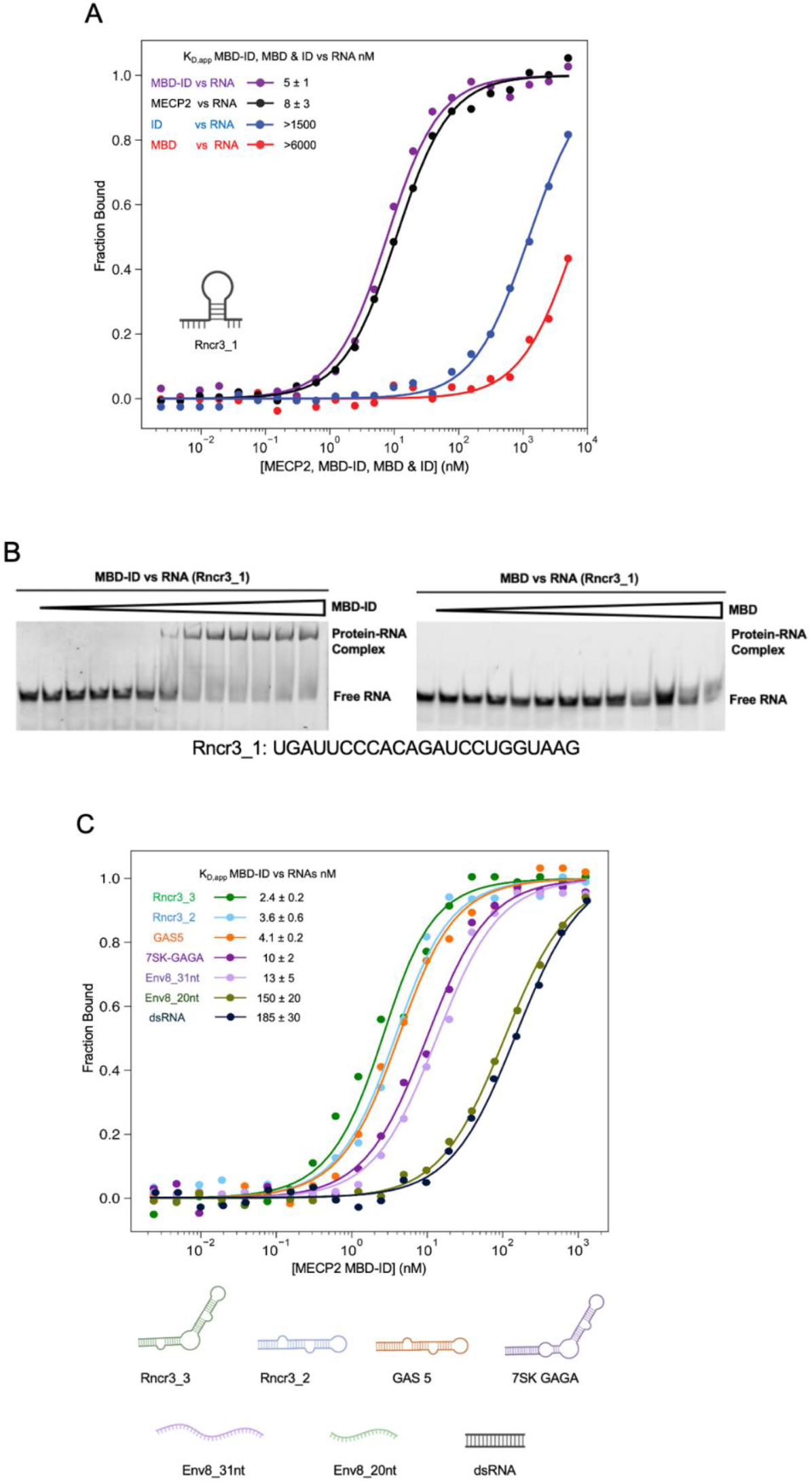
The MECP2 MBD-ID is a high-affinity, promiscuous RNA-binding module. (A) FA binding data of FL MECP2, MBD-ID, MBD and, ID bound to Rncr3_1 RNA. (B) Quantitative EMSA of MBD-ID and MBD bound to Rncr3_1 RNA. RNA concentration = 10 nM. Highest protein concentration = 1 μM, protein concentration decreases by a factor of 2 in lanes to the left. (C) FA binding data of MBD-ID bound to various RNA ligands. n>3, standard error of the mean reported.

We next sought to characterize the RNA-binding landscape of the MBD-ID module. Based on our previous work demonstrating a biologically relevant interaction between MECP2 and the lncRNA *Rncr3 (13)*, we first probed two additional *Rncr3*-derived RNA segments (Rncr3_2, Rncr3_3) predicted by UNAfold (33) to form stem-loop structures. The MBD-ID construct bound these two RNA ligands with low nanomolar affinity (Fig. 4C, K_D,app_ ≈ 2-4 nM), a range comparable to its affinity for methylated DNA (Fig. 3B).

To determine if this high-affinity binding was specific to *Rncr3*, we tested structured RNA targets of the transcription factors glucocorticoid receptor (GR) (34) and GATA1 (35), including the GAS5 and 7SK-GAGA hairpins. The MBD-ID bound these RNAs with similarly tight affinity (K_D,app_ ≈ 4-10 nM) (Fig. 4C). This broad recognition of diverse structured RNAs, akin to other proteins with arginine-rich motifs (ARMs) like the estrogen receptor (ERα) (18), GR (34) and GATA1 (35), suggests that the MBD-ID lacks intrinsic sequence specificity for RNA and instead exhibits a preference for structural features.

We probed further this specificity by testing a double-stranded RNA (dsRNA) mimicking the BDNF promoter, but this was bound with only moderate affinity by MBD-ID (K_D,app_ 185 ± 30 nM) (Fig. 4C), indicating that double-stranded RNA structure is not sufficient for high-affinity binding. We tested single-stranded RNAs (ssRNAs) of varying lengths that are unlikely to form stable secondary structures. Here, we observed a clear length-dependent affinity, with MBD-ID binding a 31-nt env8 sequence approximately 11-fold more tightly than a shorter 20-nt variant (Fig. 4C). This finding is consistent with a recent SELEX-based study (32) and implies that electrostatic interactions, contributed by an extended interaction surface, are a key determinant in non-specific binding. Overall, the MBD-ID module of MECP2 is a high-affinity but low-specificity RNA-binding unit, exhibiting a pronounced preference for RNAs capable of forming secondary structures, rather than a defined primary sequence.

### RNA-Mediated Displacement of MECP2 from DNA Requires the Intervening Domain

Another outstanding question is whether DNA- and RNA-binding through the MBD-ID can occur simultaneously or whether this is a competitive interaction (12). To test this, we employed a FA competition assay by pre-forming complexes of MECP2 constructs with fluorescently labeled mBDNF and challenging the complex with a competitor nucleic acid. As a control, we first tested unlabeled mBDNF DNA and found this could efficiently compete for binding, leading to displacement of all constructs (FL MECP2, MBD-ID, and MBD) from the labeled DNA (Fig. 5A, B, and C). Strikingly, when unlabeled Rncr3_2 RNA was used as a competitor, it potently displaced both FL MECP2 and the MBD-ID from the DNA (Fig. 5A and B), but the isolated MBD was completely resistant to RNA competition (Fig. 5C), consistent with its weak affinity for RNA. These data reinforce our finding that the MBD and ID do not function as independent modules but as an integrated functional unit. The binding of MBD-ID to one nucleic acid, be it DNA or RNA, sterically or allosterically precludes its simultaneous binding to the other nucleic acid, providing biochemical limits on suggested models for RNA-binding functions.

**Figure 5:**
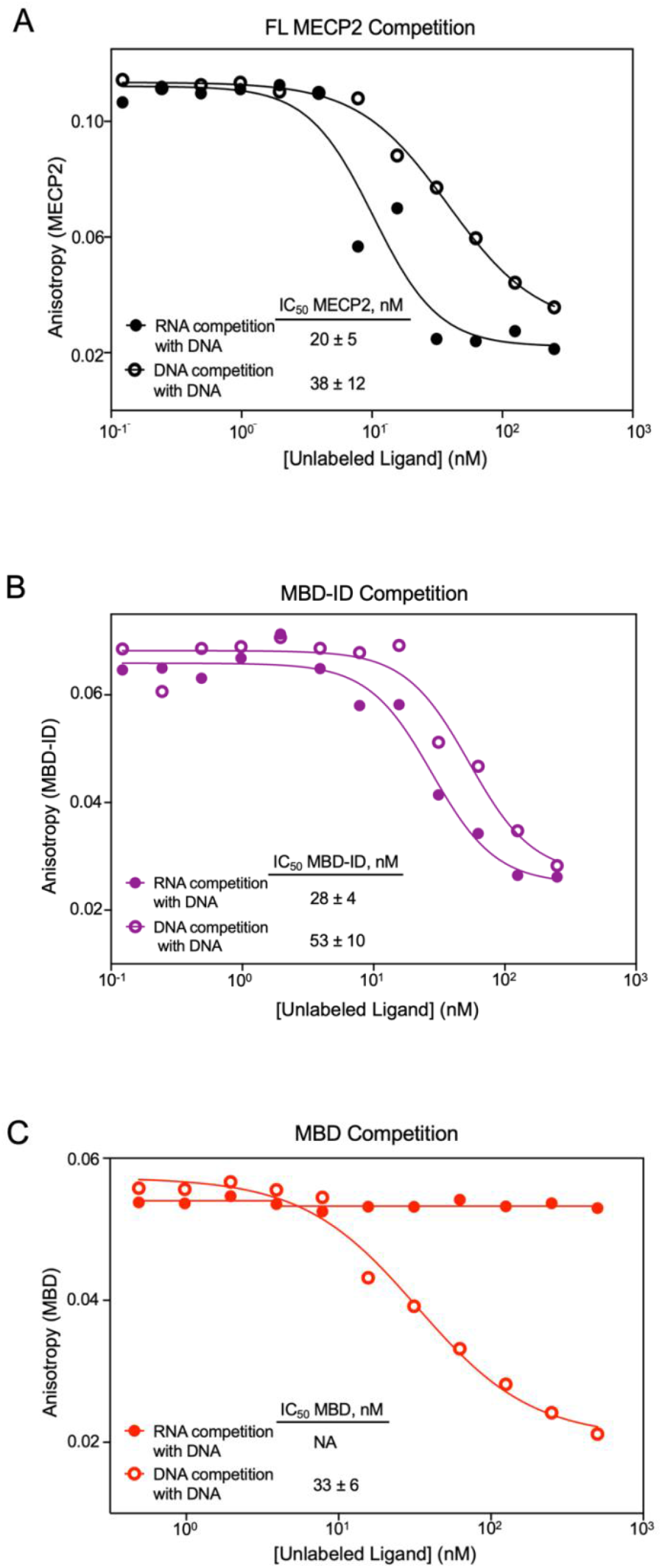
RNA-mediated displacement of MECP2 from DNA requires the intervening domain. (A) FL MECP2, (B) MBD-ID, or (C) MBD bound to labeled mBDNF DNA followed by addition of unlabeled mBDNF (hollow circles) or Rncr3_2 RNA (solid circles) at increasing concentrations. n ≥ 3, standard error of the mean reported.

### Rett-associated Mutations in the Intervening Domain Affect Nucleic Acid Interactions

The profound neurodevelopmental deficits observed in Rett syndrome (1, 20) due to *MECP2* mutations underscores the critical importance of MECP2 molecular functions. Pathogenic mutations that map to the MBD and transcription repression domain (TRD) are well studied for their impact on MECP2 biology (2). However, missense mutations within the ID are also associated with Rett syndrome and other *MECP2*-related neurodevelopmental disorders (14, 20) (Fig. 1A), although they are largely of unknown pathogenicity as little attention to date has been paid to possible ID function or its biophysical activities (2). To bridge this gap between genetics and mechanism, we quantitatively profiled the nucleic acid binding properties of three patient-derived ID variants (R167W, K174Q, and R190H) within the context of the MBD-ID module. These mutations were selected for their established clinical significance: R167W is associated with familial mental dysfunction (36), K174Q with autism spectrum disorder (14), and R190H with atypical Rett phenotype and defects in DNA binding and chromatin compaction (20). These were compared to a positive control mutant, K(171-174)A, previously shown by our lab to disrupt MECP2-*Rncr3* interaction (13). We also characterized a therapeutic candidate, the MiniGene (ΔNIC) (37), a miniaturized MECP2 variant which retains an N-terminal segment, the MBD, the first 13 amino acids of the ID, an SV40 nuclear localization signal (NLS), and the NID (Fig. 1A, B). This construct is currently in clinical trials for Rett syndrome gene therapy (FDA: TSHA 102).

All three patient variants and the K(171-174)A control bound methylated DNA (mBDNF) with an affinity nearly identical to that of wild-type MBD-ID and FL MECP2 (Table 1). This indicates that the core methyl-DNA binding function, governed primarily by the MBD, remains largely intact. However, a clear defect emerged when binding was probed to non-methylated DNA (BDNF) and, notably, to Rncr3_1 RNA. Here, all pathogenic mutants exhibited a weakened affinity, typically showing a 2-to 4-fold reduction in BDNF DNA and RNA binding compared to the FL MECP2 and MBD-ID (Table 1). These data reveal a common molecular phenotype for at least some Rett syndrome-associated ID mutations: the preservation of methyl-DNA binding, and a reproducible impairment in the recognition of non-methylated DNA and RNA. This establishes that pathogenic dysfunction can arise not only from a loss of the canonical DNA binding activity, but from a more subtle decoupling of the dual nucleic acid binding capabilities of the MBD-ID module, highlighting the essential role of the ID in fine-tuning MECP2 genomic and epitranscriptomic interactions.

**Table 1:**
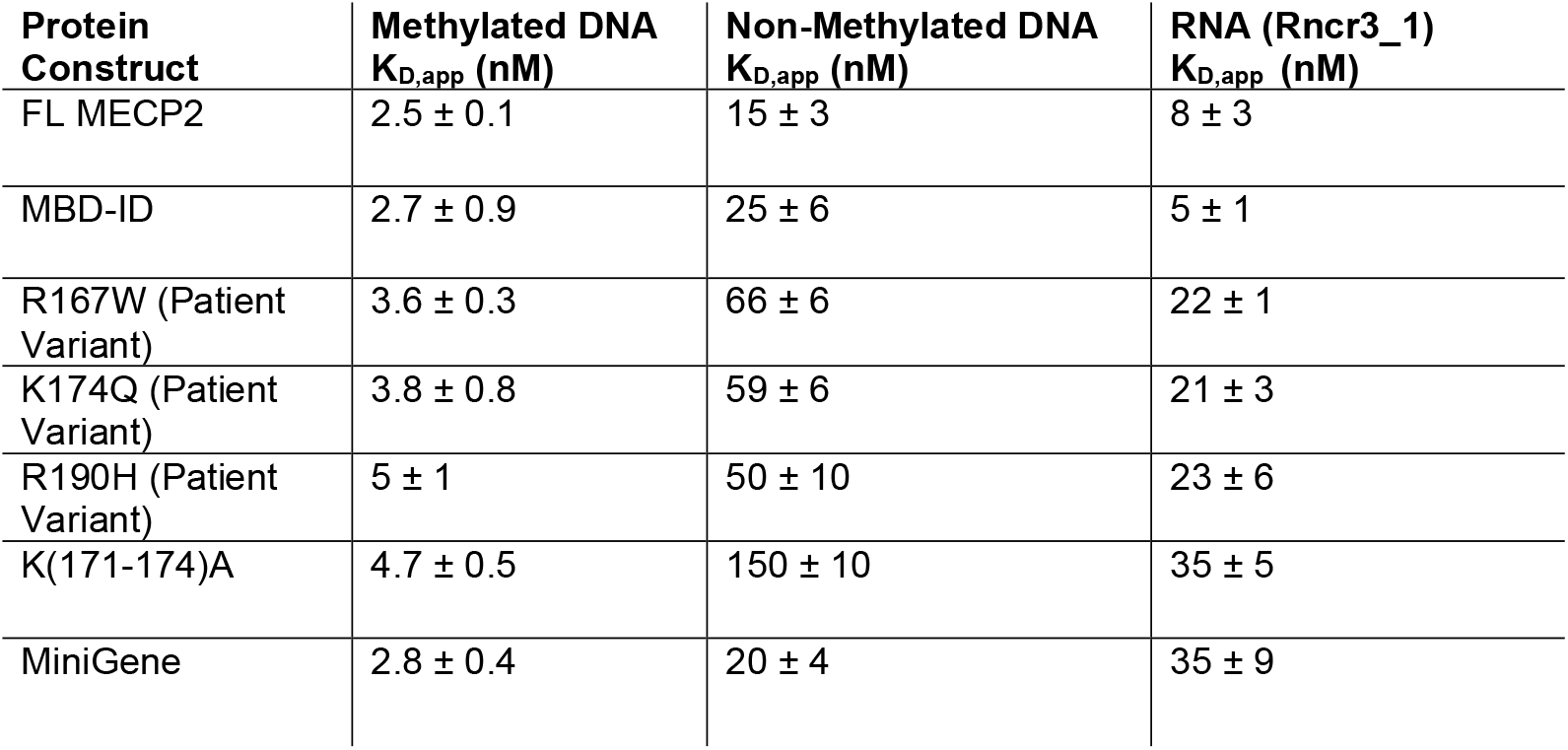
Rett-associated mutations in the intervening domain and their effect on nucleic acid interactions.

| **Protein Construct** | **Methylated DNA**<br>**$K_{D,app}$ (nM)** | **Non-Methylated DNA**<br>**$K_{D,app}$ (nM)** | **RNA (Rncr3_1)**<br>**$K_{D,app}$ (nM)** |
| --- | --- | --- | --- |
| FL MECP2 | $2.5 \pm 0.1$ | $15 \pm 3$ | $8 \pm 3$ |
| MBD-ID | $2.7 \pm 0.9$ | $25 \pm 6$ | $5 \pm 1$ |
| R167W (Patient Variant) | $3.6 \pm 0.3$ | $66 \pm 6$ | $22 \pm 1$ |
| K174Q (Patient Variant) | $3.8 \pm 0.8$ | $59 \pm 6$ | $21 \pm 3$ |
| R190H (Patient Variant) | $5 \pm 1$ | $50 \pm 10$ | $23 \pm 6$ |
| K(171-174)A | $4.7 \pm 0.5$ | $150 \pm 10$ | $35 \pm 5$ |
| MiniGene | $2.8 \pm 0.4$ | $20 \pm 4$ | $35 \pm 9$ |

Having established the role of the ID in a unified DNA/RNA binding interface, we next asked how this function is represented in the therapeutic MiniGene. The profile of the MiniGene was particularly informative. It recapitulated wild-type-level binding to both methylated and non-methylated DNA. However, its affinity for RNA was reduced more than 5-fold relative to FL MECP2 and MBD-ID, a defect more compromised than the patient variants. This biochemical signature suggests that while the MiniGene retains robust DNA binding capacity, its interaction with the RNA interactome is altered, underscoring the need to consider the engineering of optimized therapeutic protein constructs in the context of their biochemical properties.

## DISCUSSION

Our study redefines the mechanistic paradigm of MECP2 by demonstrating that the methyl-CpG- and RNA-binding activities are not segregated but are integrated functions of a single, synergistic MBD-ID module contained within the full-length protein. This model resolves the conflicting evidence (12) of how RNA can displace MECP2 from DNA: it is not a competition between independent domains, but a competition for a functionally overlapping interface. The synergy we observe, with the ID enhancing the MBD’s DNA affinity by 35-fold and RNA affinity by over three orders of magnitude, underscores that the MBD and ID are co-dependent elements necessary and sufficient for the nucleic acid binding of MECP2. This has direct and critical implications for understanding the effect of mutations that lie within the highly conserved ID (Fig. 1), many of which are classified as variants of unknown significance (14). We found that pathogenic mutations within the ID lead to a molecular decoupling, that is, they preserve high-affinity methyl-DNA binding while selectively impairing interactions with non-methylated DNA and RNA. The MiniGene, which is being tested in the clinic (FDA: TSHA 102), preserves DNA-binding functions but exhibits diminished RNA binding. Modulating this balance of nucleic acid binding activities may have better therapeutic outcomes.

We speculate that the physiological function of this dynamic, competitive interplay between DNA- and RNA-binding is to modulate MECP2 chromatin occupancy. Given the exceptionally high levels of MECP2 in mature neurons (38) and the abundance of its methyl-CpG/CpH recognition sites (39), a mechanism must exist to prevent irreversible chromatin condensation and allow for dynamic gene regulation. Our data points to a potential model where RNA binding could serve as a critical rheostat for MECP2 chromatin occupancy. Nuclear RNAs, including nascent transcripts and specific lncRNAs, could act as molecular competitors that can actively displace a subset of MECP2 molecules from chromatin to fine-tune gene regulation. A critical future step to test this model will be determining the structural basis of this integrated binding. High-resolution structures of the MBD-ID module bound to methylated DNA and to RNA would reveal the precise molecular contacts that define the competitive interface. This structural knowledge is essential for the rational design of separation-of-function mutations that selectively disrupt RNA binding while preserving DNA binding. These engineered variants would then serve as powerful tools to dissect the potential contributions of MECP2-RNA interactome to chromatin organization and transcriptional regulation in neurons.

This potential model positions MECP2 at a nexus: it is a chromatin organizer whose own placement is regulated by the epitranscriptome. The pathology of *MECP2*-related neurological disorders may therefore arise not only from disrupted DNA binding and chromatin architecture but also from dysfunctional RNA interaction. Future studies should help to reveal whether ID mutations and disrupted RNA interaction cause hyperstability of MECP2 on chromatin, leading to reduced ability to dynamically modulate gene expression during neural cell fate transitions and establishment of functional neural circuits. Together our studies suggest that the MBD-ID module acts as a molecular processor of nucleic acid cues, and its balanced output is essential for neurodevelopment.

## ACKNOWLEDGEMENTS

We thank Dr. Samuel Hunter for bioinformatics analysis and feedback on manuscript. We are grateful to Chelsea Drown for molecular cloning and to Daniella Ugay for technical support and critical feedback. We thank Dr. Vignesh Kasinath for providing the full-length MECP2 construct and for advice on protein purification. We also acknowledge the use of shared instrumentation supported by the NIH Shared Instrumentation Grant S10OD034218-0 in the Department of Biochemistry, University of Colorado Boulder.

## AUTHOR CONTRIBUTIONS

**J.A.P:** Conceptualization, Investigation, Formal Analysis, Visualization, Writing (Original draft, review and editing)

**T.A.W:** Protein Biochemistry (Methodology and Resource)

**R.T.B:** Resources, Supervision, Writing Reviews and, Funding Acquisition

**L.A.N:** Conceptualization, Supervision, Funding Acquisition, Project Administration, Writing Review.

**D.S.W:** Resources, Supervision, Project Administration, Writing Reviews and Funding Acquisition

## SUPPLEMENTARY DATA

Supplementary Data are available online.

## CONFLICY OF INTEREST

The authors declare no competing interests.

## FUNDING

This work was supported by an Innovation Award from the International Rett Syndrome Foundation and by NIH NINDS R01NS110887 award (to L.A.N), RO1GM120347 (to D.S.W) and R35 GM152029 (to R.T.B.).

## DATA AVAILABILITY

## Notes

### Competing Interest Statement

The authors have declared no competing interest.

