## Supplementary Data for "MECP2 MBD-ID Module: A Unified DNA/RNA Binding Interface Disrupted in Rett Syndrome"

**Table S1. Recombinant Protein sequence used in this study.**

| Name | Residue | Affinity Tag & Protease Recognition sequence | Amino Acid Sequence |
|---|---|---|---|
| FL MECP2 | 1-486 | Strep-Tag II WSHPQFEKGA ENLYFQSNA | MVAGMLGLREEKSEDQDLQGLKDKPLKFKKVKKDK KEEKEGKHEPVQPSAHHSAEPAEAGKAETSEGSGS APAVPEASASPKQRRSIIRDRGPMYDDPTLPEGWTR KLKQRKSGRSAGKYDVYLINPQGKAFRSKVELIAYFE KVGDTSLDPNDFDFTVTGRGSPSRREQKPPKKPKS PKAPGTGRGRGRPKGSGTTRPKAATSEGVQVKRVL EKSPGKLLVKMPFQTSPGGKAEGGGATTSTQVMVIK RPGRKRKAEADPQAIPKKRGRKPGSVVAAAAAEAKK KAVKESSIRSVQETVLPIKKRKTRETVSIEVKEVVKPL LVSTLGEKSGKGLKTCKSPGRKSKESSPKGRSSSAS SPPKKEHHHHHHHSESPKAPVPLLPPLPPPPPEPES SEDPTSPPEPQDLSSSVCKEEKMPRGGSLESDGCP KEPAKTQPAVATAATAAEKYKHRGEGERKDIVSSSMP RPNREEPVDSRTPVTERVS |
| MBD | 78-165 | Hexahistidine MGSSHHHHH HSSENLYFQS H | SASPKQRRSIIRDRGPMYDDPTLPEGWTRKLKQRKS GRSAGKYDVYLINPQGKAFRSKVELIAYFEKVGDTSL DPNDFDFTVTGRGSP |
| MBD-ID | 78-207 | Hexahistidine MGSSHHHHH HSSENLYFQS H | SASPKQRRSIIRDRGPMYDDPTLPEGWTRKLKQRKS GRSAGKYDVYLINPQGKAFRSKVELIAYFEKVGDTSL DPNDFDFTVTGRGSPSRREQKPPKKPKSPKAPGTG RGRGRPKGSGTTRPKAATSEGV |
| MBD-ID R168W | 78-207 | Hexahistidine MGSSHHHHH HSSENLYFQS H | SASPKQRRSIIRDRGPMYDDPTLPEGWTRKLKQRKS GRSAGKYDVYLINPQGKAFRSKVELIAYFEKVGDTSL DPNDFDFTVTGRGSPSRWEQKPPKKPKSPKAPGTG RGRGRPKGSGTTRPKAATSEGV |
| MBD-ID K174Q | 78-207 | Hexahistidine MGSSHHHHH HSSENLYFQS H | SASPKQRRSIIRDRGPMYDDPTLPEGWTRKLKQRKS GRSAGKYDVYLINPQGKAFRSKVELIAYFEKVGDTSL DPNDFDFTVTGRGSPSRREQKPPQKPKSPKAPGTG RGRGRPKGSGTTRPKAATSEGV |
| MBD-ID R190H | 78-207 | Hexahistidine MGSSHHHHH HSSENLYFQS H | SASPKQRRSIIRDRGPMYDDPTLPEGWTRKLKQRKS GRSAGKYDVYLINPQGKAFRSKVELIAYFEKVGDTSL DPNDFDFTVTGRGSPSRREQKPPKKPKSPKAPGTG RGRGHPKGSGTTRPKAATSEGV |
| MBD-ID K(171-174)A | 78-207 | Hexahistidine MGSSHHHHH HSSENLYFQS H | SASPKQRRSIIRDRGPMYDDPTLPEGWTRKLKQRKS GRSAGKYDVYLINPQGKAFRSKVELIAYFEKVGDTSL DPNDFDFTVTGRGSPSRREQAPPAAPKSPKAPGTG RGRGRPKGSGTTRPKAATSEGV |
| MBP-ID | ID (161-207) | Octahistidine MHHHHHHHH | MHHHHHHHHKIEEGKLVIWINGDKGYNGLAEVGKKF EKDTGIKVTVEHPDKLEEKFPQVAATGDGPDIIFWAH DRFGGYAQSGLLAEITPDKAFQDKLYPFTWDAVRYN |

| | | | |
|---|---|---|---|
| | | | GKLIAYPIAVEALSLIYNKDLLPNPPKTWEEIPALDKEL KAKGKSALMFNLQEPYFTWPLIAADGGYAFKYENGK YDIKDVGVDNAGAKAGLTFLVDLIKNKHMNADTDYSI AEAAFNKGETAMTINGPWAWSNIDTSKVNYGVTVLP TFKGQPSKPFVGVLSAGINAASPNKELAKEFLENYLL TDEGLEAVNKDKPLGAVALKSYEEELAKDPRIAATME NAQKGEIMPNIPQMSAFWYAVRTAVINAASGRQTVD EALKDAQTNSSSVPGRGSIEGR GRGSPSRREQKPPKKPKSPKAPGTGRGRGRPKGS GTTRPKAATSEGV |
| MBP | | Octahistidine MHHHHHHHH | MHHHHHHHHKIEEGKLVIWINGDKGYNGLAEVGKKF EKDTGIKVTVEHPDKLEEKFPQVAATGDGPDIIFWAH DRFGGYAQSGLLAEITPDKAFQDKLYPFTWDAVRYN GKLIAYPIAVEALSLIYNKDLLPNPPKTWEEIPALDKEL KAKGKSALMFNLQEPYFTWPLIAADGGYAFKYENGK YDIKDVGVDNAGAKAGLTFLVDLIKNKHMNADTDYSI AEAAFNKGETAMTINGPWAWSNIDTSKVNYGVTVLP TFKGQPSKPFVGVLSAGINAASPNKELAKEFLENYLL TDEGLEAVNKDKPLGAVALKSYEEELAKDPRIAATME NAQKGEIMPNIPQMSAFWYAVRTAVINAASGRQTVD EALKDAQTNSSSVPGRGSIEGR |
| $h\Delta$NIC MiniGene | | Hexahistidine MGSSHHHHH HSSENLYFQS H | MVAGMLGLREEKPAVPEASASPKQRRSIIRDRGPMY DDPTLPEGWTRKLKQRKSGRSAGKYDVYLINPQGK AFRSKVELIAYFEKVGDTSLDPNDFDFTVTGRGSPSR REQKPPGSSGSSGPKKKRKVPGSVVAAAAAEAKKK AVKESSIRSVHETVLPIKKRKTRETV |

Key: Amino acid in red represent Rett patient ID mutation
In the MBP-ID sequence, amino acid in blue represents the ID sequence.

**Table S2. Nucleic Acid Oligonucleotides used in this study.**

| Feature | Name | Sequence 5' – 3' | Label | Size |
|---|---|---|---|---|
| dsDNA | mBDNF_Top | CTGGAAmCGGAATTCT | 5'-6FAM | 15 nt |
| dsDNA | mBDNF_Bot | AGAATTCmCGTTCCAG | | 15 nt |
| dsDNA | BDNF_Top | CTGGAACGGAATTCT | 5'-6FAM | 15 nt |
| dsDNA | BDNF_Bot | AGAATTCCGTTCCAG | | 15 nt |
| ssRNA | Env8_20nt | AUACAACAUACAACAUACAA | 5'-6FAM | 20 nt |
| ssRNA | Env8_31nt | AUACAACAUACAACAUACAACAUA CAACAUC | 5'-6FAM | 31 nt |
| dsRNA | BDNF RNA_Top | CUGGAACGGAAUUCU | 5'-6FAM | 15 nt |
| dsRNA | BDNF RNA_Bot | AGAAUUCCGUUCCAG | | 15 nt |
| ssRNA | Rncr3_1 | UGAUUCCCACAGAUCCUGGUAAG | 5'-6FAM | 23 nt |
| ssRNA | Rncr3_2 | GGGGAGAGUUCCGGAAGGCUGA UUCCCCC | 5'-6FAM | 29 nt |
| ssRNA | Rncr3_3 | GGCCUGGUAAGGGACCGCAGCA GCUCUGCCCUGCGGCACGCCCG GCCAGGCC | 5'-6FAM | 52 nt |
| ssRNA | Gas5 | GGGAGCCUCCCAGUGGUCUUUG UAGACUGCCUGAUGGAGUCUCC CC | 3'-AF488 | 46 nt |
| ssRNA | 7SK-GAGA | GGGAUCUGUCACCCCAUUGAUC GCCGAGAGGCUGAUCUGGCUGG CUAGGCGGGUCCCC | 3'-AF488 | 58 nt |

Key: mC = methylated cytosine

**Table S3: Primers used to generate Rett syndrome patient ID mutants and control**

| Template | Forward Primer 5'-3' | Reverse Primer 5'-3' | Target construct |
|---|---|---|---|
| MBD-ID | CTGCGGCGCCCAAATCTCCCAAAGCTC | GTGGCGCCTGCTCTCGCCGGG | K(171-175)A |
| MBD-ID | GAGCCCCTCCTGGCGAGAGCAGA | CCTCTCCCAGTTACCGTGAAGTCAAAATCATTAG | R167W |
| MBD-ID | GAAACCACCTCAGAAGCCCAAATC | TGCTCTCGCCGGGAG | K174Q |
| MBD-ID | AGGCCGGGGACATCCCAAAGGGA | CTGCCAGTTCCTGGAGC | R190H |

| Lanes | 1 | 2 | 3 | 4 | 5 | 6 |
|---|---|---|---|---|---|---|
| Sample | Ladder | WT-MBD-ID | MBD-ID K(171-175)A | R167W | K174Q | R190H |
| Mol. Wt. KDa | 10-180 | 14 | 14 | 14 | 14 | 14 |

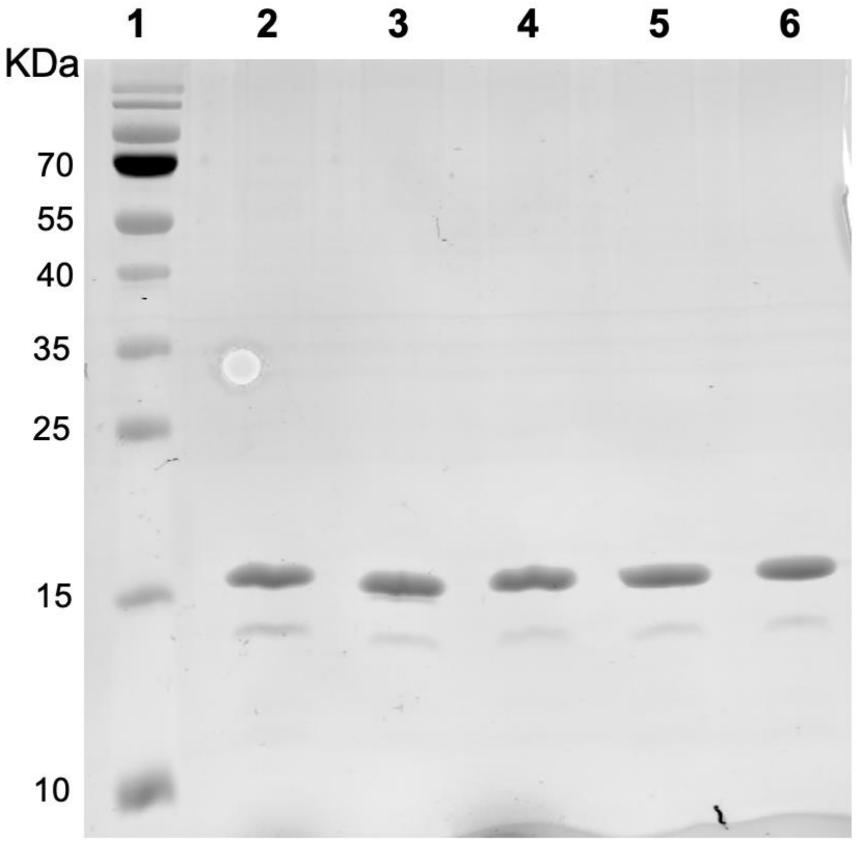


**Figure S1**. AcquaStain stained SDS PAGE gel (15% acrylamide/bis) of the purified recombinant Rett syndrome patient ID mutant protein, and control protein used in this study. Gel imaged on Typhoon FLA 9500 Imager (GE Healthcare) using Coomassie green filter.

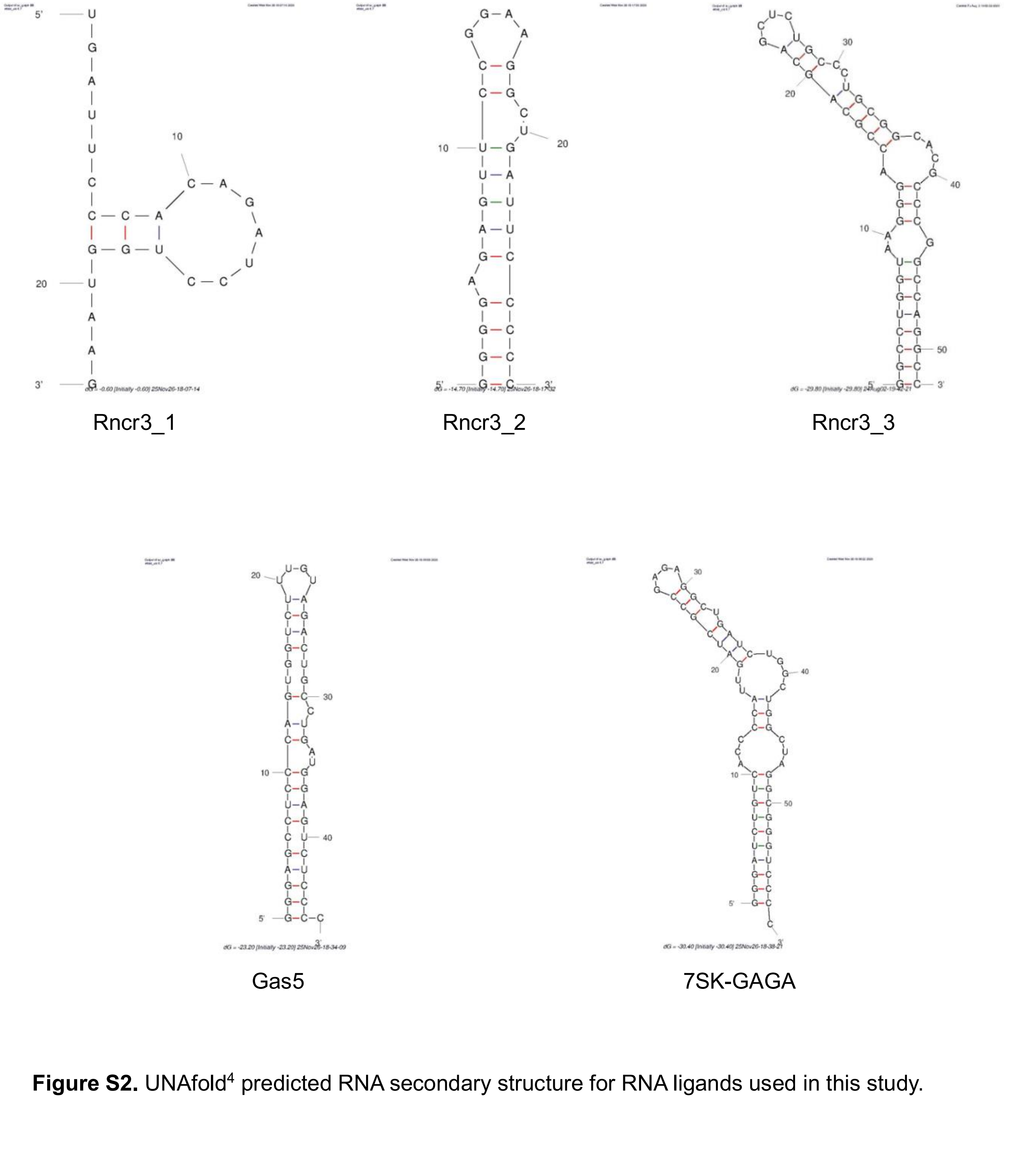


**Figure S2.** UNAfold[4] predicted RNA secondary structure for RNA ligands used in this study.

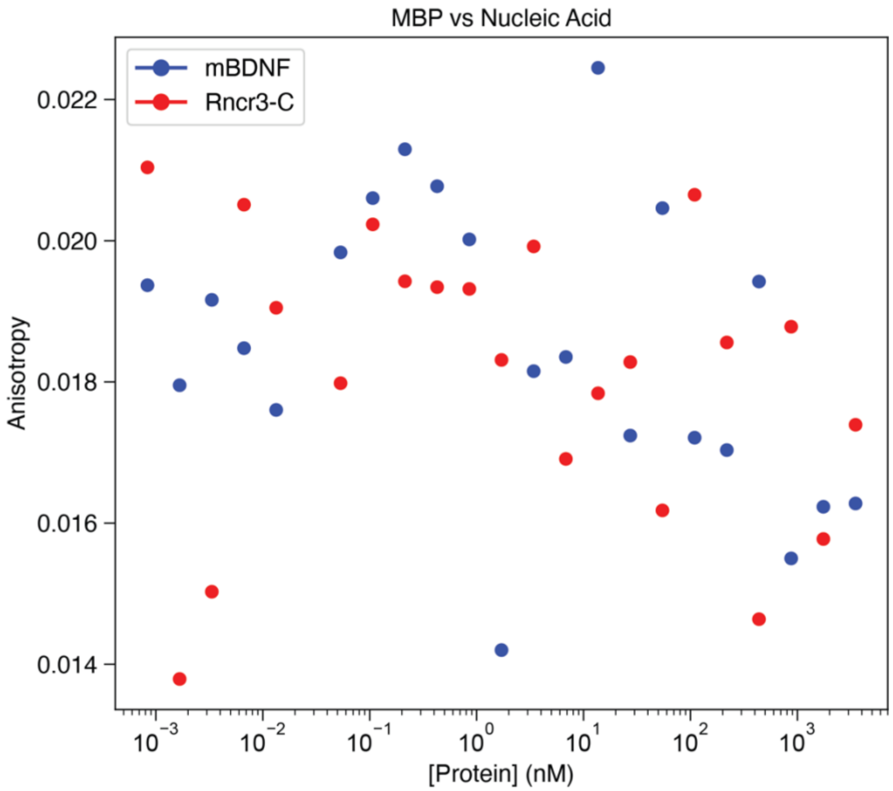


**Figure S3.** Isolated MBP does not interact with nucleic acid. Fluorescent anisotropy (FA) binding data for MBP bound to methylated BDNF DNA (blue) and Rncr3_1 RNA (Red). n>3.
